# Topology highlights mesoscopic functional equivalence between imagery and perception

**DOI:** 10.1101/268383

**Authors:** Esther Ibáñez-Marcelo, Lisa Campioni, Angkoon Phinyomark, Giovanni Petri, Enrica L. Santarcangelo

## Abstract

The functional equivalence between mental images and perception or motion has been proposed on the basis of neuroimaging evidence of large spatially overlapping activations between real and imagined sensori-motor conditions. However, similar local activation patterns do not imply the same mesoscopic integration of brain regions active during imagery and perception or action. Here we present the first EEG evidence of topological equivalence between functional network organization at intermediate and global scales during tasks. We show that the degree of functional equivalence varies in the population and is associated with different magnitudes in the restructuring of the functional connectivity between imagery and real tasks. In particular, changes observed during imagery with respect to basal conditions account for the cognitive effort experienced during imagery, and subjects characterized by stronger functional equivalence exhibit smaller topological deviations in the imagination tasks performed after real tasks, thus showing learning effects. Altogether, our findings point to different sensori-cognitive information processing in the subjects showing different functional equivalence. We anticipate our results to be a starting point for a novel dynamical description of functional equivalence, which will be relevant for socio-cognitive theories of embodiment and cognitive formulations of how different selves emerge from neurophysiological assets.

---

Mental imagery is one of our most striking cognitive capacities. It can be described as “the ability to generate, represent and manipulate objects and events that are not physically present”^1^. It has long attracted intense research as it plays an important role in human cognition, being involved in memory, abstract/spatial reasoning, action planning, decision making, skill learning and language comprehension^2, 3^. Researchers have long debated which cognitive process allows us to form perceptual/motor mental images when the corresponding sensory stimuli are absent^4^. The leading hypothesis, formalized by Finke^5^ and leading back to James^6^, is that a substantial “functional equivalence (FE)” exists between imagery and perception (or motion). That is, the two cognitive activities share the same neurophysiological bases, and imagery is an actual physical simulation of perceptual and motor experience^7–9^. Evidence of shared mechanisms has been provided mainly by neuroimaging studies which have shown partially similar brain activations during action or perception of all sensory modalities and the correspondent mental images^2, 3, 10–20^. Indeed, spatial overlap between activations provides insights into which regions perform similar roles during mental images and perception. However, local information about the spatial activation is not sufficient to conclude that the temporal activation profiles evolve similarly. In fact, while the same regions might activate in perception and mental images, their respective activation profiles might be differently correlated^21^, and therefore activations alone are not sufficient to conclude that the brain codes in the same way for sensory modalities and for mental images^22^. A stronger marker of genuine FE should therefore lie in the coherent changes of the correlation patterns among regions’ activities.

Here we investigate this hypothesis in electroencephalogram (EEG) signals recorded during real and imagined specific sensory conditions. This choice of recording modality highlights the focus on the temporal features of brain activity rather than the spatial overlap. Since individual differences in the ability of mental imagery modulate the degree of FE^12, 23–27^, we select a population for which behavioral studies have suggested a higher variability in FE^28^, that is the subjects with high scores of hypnotizability (*Highs*) compared to low hypnotizable individuals (*Lows*). Indeed, hypnotizability is a cognitive trait predicting the ability to modify perception, memory and behaviour according to specific suggestions^29^, both under hypnosis and in the ordinary state of consciousness^30^. Suggestions are instructions to imagine sensory-cognitive contexts different from the real one, as occurs for the widely known suggestions of analgesia that allow to control pain^31^. Hypnotizability is measured by scales^32^ that classify high, low and medium hypnotizable subjects (respectively Highs, Lows and Mediums) who often exhibit characteristics intermediate between Highs and Lows^33^. Different levels of hypnotizability have been associated with structural and functional brain peculiarities^34–36^, differences in sensorimotor integration and cardiovascular control^37^ as well as in imagery abilities^38–42^ and in the behavioral effects of imagery^43–48^. In particular, behavioural studies of the vestibulo-spinal reflex (VSR) suggested stronger FE in Highs^48^. The VSR, elicited by galvanic stimulation of the labyrinth, induces body sway mainly in the frontal plane when the head is directed forward and in the sagittal plane when the head is rotated toward one side^49^. Importantly, the VSR earlier component cannot be voluntarily modulated^50, 51^. In the above cited behavioural study^48^ Highs and Lows were guided to imagine the rotated position of the head through both the visual and the kinestethic sensory modality. Interestingly, despite the similar vividness of the mental image of rotated head, Highs, but not Lows, exhibited the same VSR earlier component during the real and the imagined position of rotated head. On these bases, we chose to study FE in Highs and Lows because they are expected to exhibit very different degrees of FE.

We analyse here the correlation patterns of EEG signals from a topological point of view^52^. Topology^53^, one of the deepest branches of mathematics, describes the shape and connectivity of spaces in any dimension. Tools based on topological ideas, collectively dubbed Topological Data Analysis (TDA)^54^, emerged over the last decade to study systems where relevant scales range between different spatial and temporal resolutions, and information is encoded in higher-order interactions^55^. They provide a natural language to describe local, mesoscopic and global features of data, and are therefore well-suited to capture the mesoscale functional organization of activation patterns. Indeed, the shape and the topological features of a dataset –be it an image, a network or a matrix– is effectively described by the patterns of dense areas and the regions of disconnectivity among the former. The structure and properties of these holes define–in a Spinozian way– the structure of the whole dataset. This is called the homology of the dataset, and its graded version, persistent homology, is able to deal with weighted, noisy, un-evenly sampled and complex datasets. It works by producing a sequence of progressively finer approximations –called *filtration*– of the data and by tracking the evolution of the voids across such filtration. Important features and structures in the data live longer through the filtration spanning a large range of approximations or multiple scales (we provide further details in the Methods and the SI).

Persistent homology techniques have been recently applied to neuroimaging data to characterize resting states of consciousness, physiological^56^, altered^57^, and dynamical^58^, and sensory tasks^59^. They have also been applied to EEG data in humans to characterize cognitive tasks^60^ and in animal models to classify normal^61^ and pathological behaviour^62^. Interestingly, topological features were also able to detect the encoding of geometric structure from neural activity alone in rat neuron spike data^63, 64^, and have been used to inform the mapping between structural and functional connectomes^65, 66^.

In light of previous findings ^48^, we hypothesize that stronger FE should be associated with vivid and effortless mental images. We show here that is indeed the case, by reporting the first topological evidence of such stronger (weaker) FE in Highs (Lows). In particular, we show that the *topological deviation* between basal and imagery states predicts the cognitive effort required by a given imagery task, and that smaller topological deviations between the imagery state and the target real state correspond to increased vividness of the mental imagery. Finally, we show that Highs exhibit reduced topological deviations in the imagination tasks after performing the real task, pointing to a more efficient sensori-cognitive information processing scheme with respect to Lows.

## Results

### Deviations from basal state during imagery predict cognitive effort

We analysed EEG signals (32 channels per subject) recorded from 18 Highs and 19 Lows. Each subject performed 5 different tasks, with each trial including a basal and task condition lasting 1 minute each. In basal conditions participants were invited to relax at their best. Visual (v) and kinestethetic (k) imagery tasks were performed both before (Time 1: v1,k1) and after (Time 2: v2, k2) a condition of real rotated position of the head (rr) in order to study potential learning effects. The order of the visual and kinesthetic imagery-which for individual subjects was the same for trials performed before and after rr - was randomized among subjects. Full details are provided in the Methods section. After each imagery task, subjects were invited to score the vividness of their mental images and the experienced cognitive effort on a scale from 0 (minimum) to 10 (maximum). For each subject, task and corresponding basal period, we computed the correlation matrices between the recorded EEG signals. We then computed the persistent homology, as described in^57, 67^ for each combination. The main output of persistent homology are *persistence diagrams* (Figure 1a): for each dimensionality of holes (zero-dimensional connected components, one-dimensional loops, cavities and higher dimension analogues), a persistence diagram is a multiset of points in two dimensions, where each point represents a specific topological hole and is labeled by its birth time across the filtration and death time. Features are generally considered more important the larger their *persistence* is. Crucially, it is possible to compute a similarity metric between persistence diagrams. The similarity of persistence diagrams is calculated using the persistence scale-space kernel proposed by^68^. This kernel is parametrized by a scale parameter *σ* which captures how wide the Gaussian kernels on each of the points in the persistence diagrams are, and hence implicitly defines how finely or coarsely we are comparing the two persistence diagrams (Figure 1b and Methods for details).

**Figure 1.**
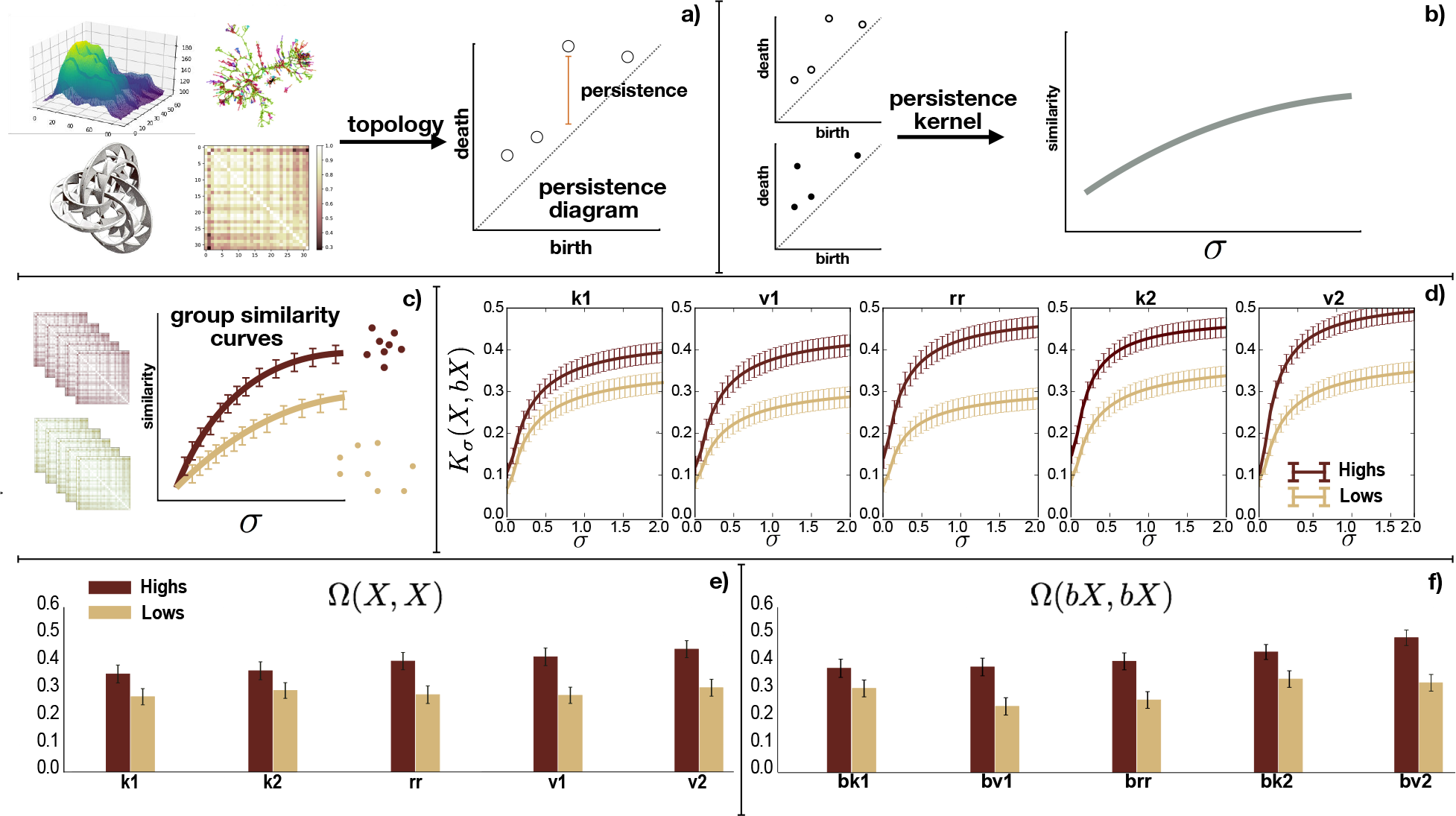
Tasks are split in visual (v) and kinesthetic (k) and *k*1, *v*1 and *k*2, *v*2 refers to tasks before and after real rotation (*rr*) respectively. a) From correlation matrix (left) to persistence diagrams (right) through persistent homology (topological features). b) Given two persistence diagrams (PDs) we define a similarity measure (depending on sigma) between them. c) Similarity measure applied on a set of PDs for two different groups. d) Comparison of similarity between Highs and Lows for all basal-task pairs *bX* → *X*: the basal deviation of Highs is always smaller (*K*_*σ*_ (*X*, *bX*) is higher) than in Lows. Confidence intervals have been computed assuming normal distribution of the sampled mean at 95% of confidence. e) and f) Higher homogeneity in Highs group respect to Lows both in basal and task condition, respectively. Confidence intervals have been computed using the bootstraping technique on the subsampled averaged on a 95% level.

A high similarity between two persistence diagrams corresponds to small differences in the topology of the two corresponding spaces. Furthermore, for a group of subjects, we can measure the similarity between all pairs of subjects within the group and compare it with the similarity within another group. These are the two lines in Figure 1c. A higher group similarity will correspond to the (homological structure of the) subjects being more uniform as group (dark brown dots) as opposed to a more heterogeneous group (light yellow dots).

First, we compare how much the subjects’ one-dimensional homological structure (i.e. the *H*_1_ group composed by cycles) changed between a task *X* and its corresponding basal *bX* (*bX* → *X*). For clarity, we dub this change the topological *basal deviation* for task *X*. The task and basal sets are respectively {*X*} = {*k*1, *v*1, *rr*, *k*2, *v*2} and {*bX*} = {*bk*1, *bv*1, *brr*, *bk*2, *bv*2}. For a subject i and a range of s values, we compute the kernel *K*_σ_ (*X*^*i*^, *bX*^*i*^) between the 1-dimensional persistence diagrams obtained for each task *X* and its basal *bX*. We then aggregate the similarities within a group (Highs, Lows) and plot them as a function of σ in Figure 1d). Remarkably, we find that, for all values of σ, Highs display a larger group-similarity than Lows. This implies that functionally Highs are more similar between basal and task conditions than it is in the case of Lows. In other words, Highs require smaller basal deviation to perform tasks.

Note that the basal deviation quantifies the functional reconfiguration needed to perform the task and can thus be considered as a task activation cost. We expect therefore that higher basal deviations should correspond to increased effort. This is consistent with subjective reports, that indicate that over all tasks Highs require lower cognitive effort than Lows (*F*(1, 35) = 4.494, *p* < .041, η^2^ = .114, see Figure S.6 in SI). Although subjective experience can rarely be reconducted to a single physiological index^69^, in this case we find further support for this interpretation from the significant anti-correlation (Spearman *R* = −0.3, *p* < 0.05) that we find between Ω(*bX*, *X*) and effort (Fig. S.7).

In addition to smaller basal deviations, Highs as a group display more homogeneous topologies. To show this, we compute the kernel similarity *K*_*σ*_ (*X*^*i*^, *X* ^*j*^) for task *X* and *K*_*σ*_ (*bX*^*i*^, *bX* ^*j*^) for the corresponding basal *bX* where we consider all pair of subjects *i*, *j* in the two groups. We did not observe any specific dependence on the value of σ, so we can eliminate it and obtain a more compact indicator of the difference between persistence diagrams by integrating away σ. Formally, for a task/basal *a*, group *g* (Highs - H, Lows - L) and subjects *i*, *j* belonging to *g*, we consider the quantities 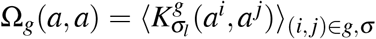, where the average is taken over all pairs and ss. In Figures 1d and 1e we report the values of Ω obtained respectively for all tasks and for all basal conditions in Highs and Lows. We find that in all conditions Highs report higher values of Ω (t-test, *p* ⪡ 0.001, *t* ∈ {4, 10}), implying that the Highs are more similar to each other than Lows are in all task and basal conditions.

### Task accuracy associates with vividness

The main claim of FE is that the neurological basis for perception and mental images is the same. Vivid perceptual mental images should therefore be associated with brain states that are close to the one attained during the actual perception. The more vivid the imagination, the more closely the imagined state should reproduce the real one, and hence the closer their topological structures should be. We study therefore the accuracy with which imagined states reproduce the target one (*rr*). We quantify the accuracy of an imaginative task *X* as Ω(*X*, *rr*). Then we ask how much imagery modifies the task patterns between Highs and Lows. We do this by considering the difference of accuracy between Highs and Lows: 
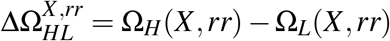

Positive values of ΔΩ_*HL*_(*X*, *rr*) mean that that the functional topologies of Highs during imagery tasks and real rotations are closer than for Lows. Note that ΔΩ is related to the area between the two similarity curves that are being compared. In Figure 2a we display the results. We find that Highs have more similar functional properties between tasks and the real rotation itself, as compared to Lows. Moreover, in the same way, we can also compare how imagery tasks are encoded before and after performing the actual head rotation, by considering 
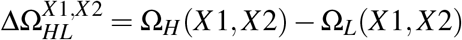

**Figure 2.**
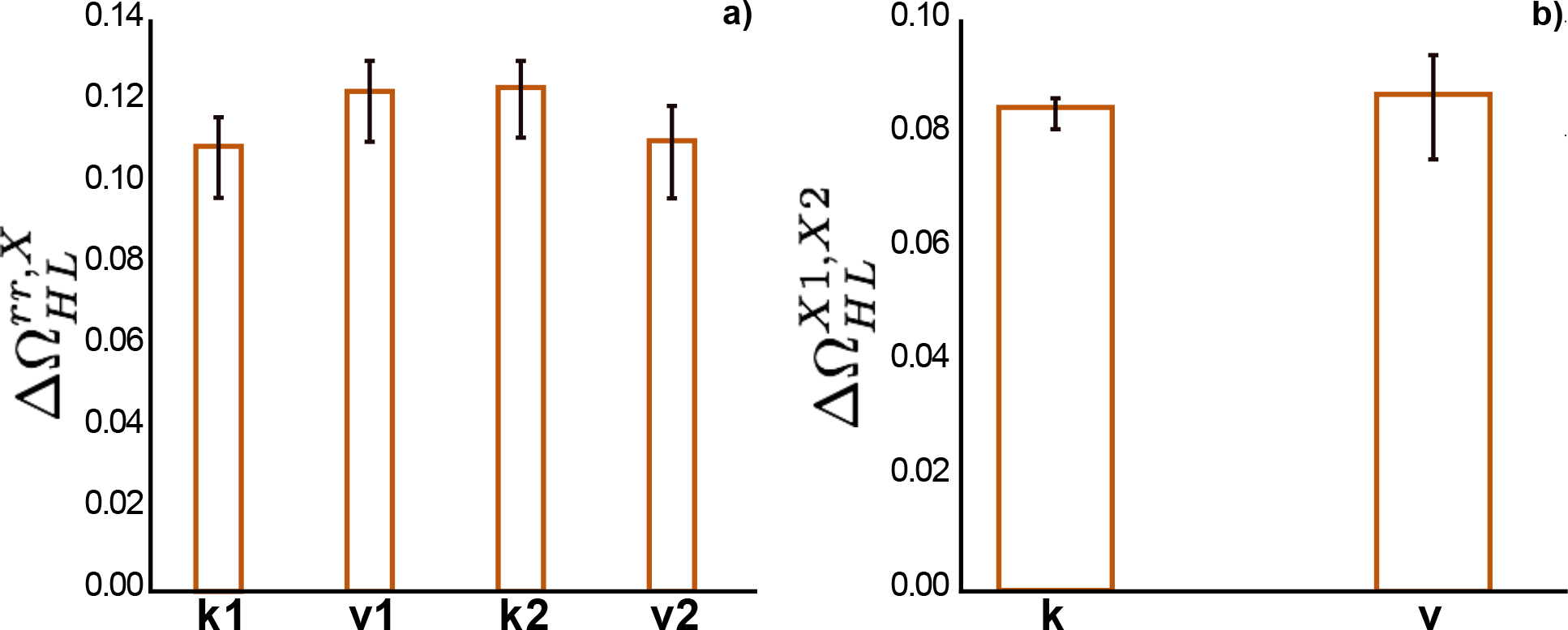
Similarity difference between imagined and actual position of rotated head. a) Difference between Highs and Lows in imagery tasks as compared to actual head rotations. Positive values indicate that Highs are topologically nearer to the real rotation than Lows. Error bars correspond to 95% confidence level on the mean. b) Difference between Highs and Lows of the alteration of topology between tasks before and after rotation (v1 vs v2, k1 vs k2). Positive results mean that Highs display more similar topologies before and after the rotation, as compared to Lows. a) & b) Confidence intervals have been computed using the bootstraping technique on the subsampled averaged on a 95% level.

We find that as a group Highs again display smaller distances between the encoding of the respective imaginations (v1/v2, k1/k2) before and after the real rotation. Consistently with the topological results, subjective reports indicate that Highs report higher vividness than Lows across both modalities (*F*(1, 35) = 11.975, *p* < .001, η^2^ = .255). Results show a significant positive difference for all imagery tasks. This indicates that –in all tasks– the topology of the imagination of Highs is nearer to that of real rotation as compared to Lows (Figure 2b) (t-test, p-val *p* < 0.05, for all values go to SI, Table S.1 and Figure S.5 in SI). With regard to the significant correlations reported between effort and basal deviation, the vividness did not significantly correlate with topological variables (Fig S.7, Spearman correlations not significant at 95%). However, the absence of significant correlation between topological and subjective indices does not challenge the relevance of the group level association accuracy-vividness, as the subjective experience is typically multidimensionally determined^69, 70^.

### Imagery performance improves in the subjects with stronger FE after real task

We showed that basal deviation is closely associated with cognitive effort, with Highs displaying small deviations and more uniformity topological structure as compared to Lows. In addition, and consistently with the vividness results, the topology of Highs’ functional structure during imagination is closer (as compared to Lows) to the one of real rotation. We ask now whether performing the real rotation before the late imagery tasks (v2,k2) induces any changes in how such imagination task is performed. From the accuracy point of view, we find for both groups negligible changes in accuracy between the early (v1,k1) and late (v2,k2) tasks. On the basis of the results of the previous section, we would therefore expect to find no significant change in the reported vividness scores. For Highs indeed we find that decomposition of the significant Time x Hypnotizability interaction (*F*(1, 35) = 5.035, *p* < .027, η^2^ = .132) reveals that Highs report the same vividness before and after the condition of real rotated head. Lows on the contrary experienced lower vividness during later tasks with respect to the earlier ones (Time, *F*(1, 18) = 4.284, *p* < .053, η^2^ = .192). Since the topological accuracy remains essentially unchanged (relative accuracy variation 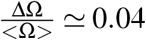, below the vividness resolution), a second factor should be responsible for the lower reported vividness in Lows. A possible explanation for this effect might lie in the different ways in which subjects perceive their own performance of the imagination. This can be coarsely quantified by comparing the basal deviation of the real rotation with that of the imagined ones. In particular, for a group g we consider 
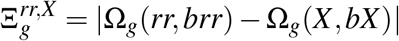

InFigure 3 we show the results for the all imaginations tasks. The difference reported point toward a substantial decrease of 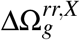 in Highs after the real rotation and an increase in Lows for both imaginations. In parallel to these results, decomposition of the significant Time × Modality × Hypnotizability interaction (*F*(1, 35) = 5.591, *p* < .024, η^2^ = .138) reveals that Highs experienced lower cognitive effort during later imagery tasks independently of the imagery modality, while Lows reported higher effort for visual imagery after the actually maintained rotation of the head than before it (Time, *F*(1, 18) = 4.788, *p* < .042, η ^2^ = .210) and no Time difference for the kinaesthetic modality.

**Figure 3.**
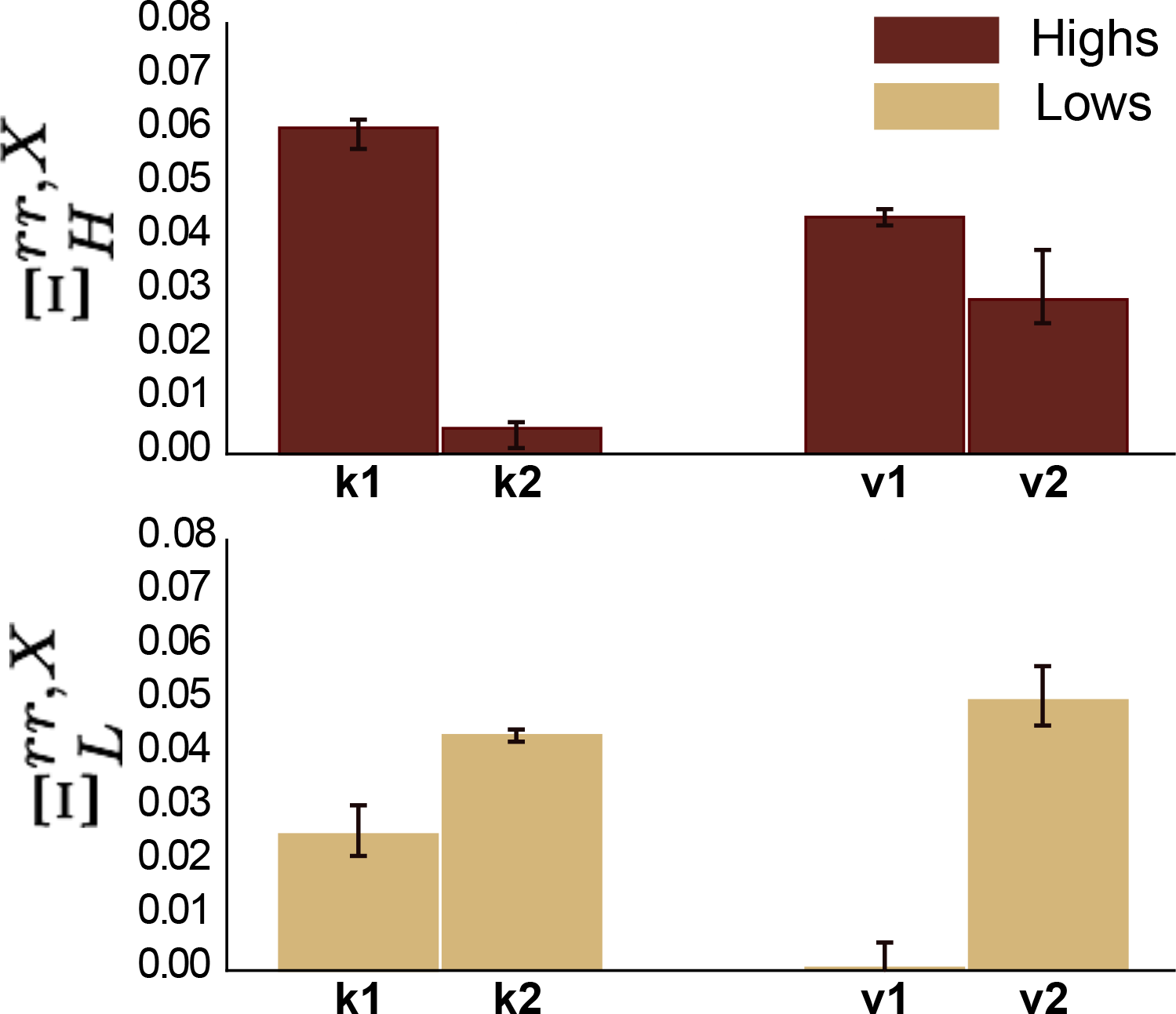
Imagery induced alteration of topology compared to rotation induced changes. We quantify using 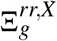 the change in effort before and after the real rotation, which can be related to task adaptation. Highs display a decrease of the difference between basal deviation between real and imagery-task, while Lows show a clear increase. This is associated with the decreased effort experienced by Highs and the increased effort experienced by Lows during later imagery tasks. Confidence intervals have been computed using the bootstraping technique on the subsampled averaged on a 95% confidence level.

We argue therefore that the reported lower vividness of Lows stems from the increased difference from the *brr* → *rr* deviation, resulting in an increased perceived effort, while not being rewarded by a significant improvement in topological accuracy. Conversely, Highs do not improve significantly their accuracy, but their reduced *brr* → *rr* distance results in lower perceived effort and improved vividness. In other words, while all subjects do not objectively improve their imagination, Highs are able to reproduce effectively the actual *brr* → *rr* transition, while Lows are not able to. These results are in line with previous evidence indicating greater imagery abilities in Highs^43, 48, 71^, which was supposed to be sustained by the larger superimposition of the activations involved in real and imagined sensory conditions in these subjects. Our results based on EEG data support this but by phrasing it in terms of similarity in co-activation shapes across different EEG signals, rather than of spatial similarity of regional activations.

## Discussion

We showed that topological properties describe hypnotizability-related differences in the functional equivalence between imagery and perception and, more broadly, in sensori-cognitive-information processing. In fact, these measures indicate that, with respect to Lows, Highs a) represent a more homogeneous sample, b) change significantly less during real and imagined perception with respect to basal conditions, c) show significantly lower differences between real and imagined perception, d) report subjective vividness of the mental images and related cognitive effort which can be accounted for by their topological features, e) display learning effects in mental imagery after experiencing real perception. Crucially, the reported differences –being topological in nature– stem from changes in the overall structure of brain activation patterns, rather than from localized or region-specific changes.

### Functional equivalence of mental images

We report the first evidence of stronger functional equivalence in Highs as they display smaller differences between the physically and imaginatively rotated head position with respect to Lows (Figure 2), supporting the hypotheses raised from behavioral studies^48^.

The stronger functional equivalence between imagery and perception observed in Highs may provide significant advancement in the physiology of the response to suggestions. In fact, theoretically a better simulation of physically induced perception may facilitate the response to sensori-motor suggestions and possibly account for the involuntariness in action reported by the Highs experiencing the subjective and behavioral effects of sensori-motor suggestions^28^.

The perception of involuntariness in action is a main component of hypnotic responding and has been differentially interpreted^72^ by the neo-dissociative^73, 74^ and socio-cognitive^75, 76^ theories of hypnosis. The former assume that hypnotic responding is reported as involuntary for the dissociation between behavior and conscious experience, which could be likely accounted for by the observed modulation of the functional connection between the salience and executive circuits in Highs^36^. In contrast, socio-cognitive theories propose that the experience of involuntariness may be sustained by peculiar configurations of cognitive, emotional, relational and sociocultural traits which make the suggested perception/behavior the most adequate to a given situation, so that it is triggered automatically and experienced as effortless and involuntary^75, 77^. We think that, at least for sensori-motor suggestions, stronger functional equivalence between imagery and perception can support the socio-cognitive views by including the degree of imagery-induced cortical activation among the many factors involved in the response to suggestions^28^.

In clinical contexts, the strength of the functional equivalence between imagery and perception may predict the efficacy of imaginative training in neuro-rehabilitation^78, 79^ and sports^9, 80^. In this respect, hypnotic assessment could become an easy and cheap tool to orient the set-up of individualized mental training.

### Sensory-cognitive information processing

We hypothesize that the scarce topological changes in Highs are the effect of largely distributed information processing. This is in line with the findings of spectral analysis revealing more widespread desynchronization in Highs than in Lows during both visual and somesthetic imagery^81^. In addition, the scarce changes elicited by tasks in Highs agree with the findings of Recurrence Quantification Analysis which failed to detect differences in the EEG recurrence plot of Highs when they did and did not receive efficacious suggestions of analgesia during nociceptive stimulation^70^.

Although the performance at cognitive and motor tasks has been associated with reconfiguration of the brain functional connectivity^82, 83^, higher intelligence has been found associated with scarce task-related brain network reconfiguration^83^ and spectral analysis supports the view that the lower the power changes the higher the cognitive performance^84–86^(Cheron et al., 2016). Indeed, we have sufficient evidence for the Highs’ extraordinary ability to perform cognitive tasks. It is indicated by their peculiar attentional ability^87, 88^, their proneness to disregard irrelevant information^89^, to voluntarily modulate their conscious experience according to specific suggestions^30^ and to modify their state of consciousness voluntarily^90^.

It has been proposed that the mechanisms underlying cognitive activities change activation patterns characterized by a more segregated, within network structure into other patterns characterized by pre-eminent between networks connectivity^91^. The shift from one to the other could be induced by ascending neuro-modulatory systems^92, 93^. The latter might be more efficient in Highs and maintain their EEG activity in a highly distributed mode of functioning independently from tasks. In Highs, in fact, we have indirect evidence of hyperactivity of the cholinergic and/or noradrenergic system indicated by their shorter reaction times in attentional tasks^94, 95^ and of higher dopaminergic tone^96^, although the mechanism classically considered responsible for it - less efficient dopamine catabolism by Catechol-O-Monoamine-Oxidase^97^ – has not been confirmed^98, 99^.

### Limitations

Although imagery is bilaterally represented in the brain^100^, the actual position of head rotated toward the right side may have activated the left and right hemisphere differentially. Thus, a limitation of the study is that the EEG laterality was not taken into account. It would have increased the number of factors of variability too much with respect to the sample size. Indeed, a much larger sample would also allow for a more sophisticated hypnotic assessment able to differentiate Highs and their topological features according to their hypnotic profile (that is the quality of the items they passed on the scoring scale) rather than simply to the total score reported on hypnotizability scales^101^. Finally, present findings allow to suggest that the Highs’ response to sensory suggestions could be facilitated by larger superimposition of the cortical activations associated with imagined and physically induced sensory experience, thus sustaining the reported involuntariness in action. Nonetheless, this mechanism may be not sufficient to explain the effects of suggestions in different domains such as distortion/suppression of memories.

## Conclusions

Topological properties provide the first evidence of EEG functional equivalence rooted in the coevolution of brain activations. They reveal hypnotizability-related sensory-cognitive information processing and of stronger functional equivalence between imagery and perception of the position of a body part in the subjects with high scores of hypnotizability. Moreover, our results provide insights of efficient versus expensive information processing between high and low type of hypnotizable subjects. Crucially, this stronger functional equivalence is supported by non-local activation patterns, rather than by local regional activations, pointing to the need to characterize the changes in functional connectivity at the mesoscopic scale^102, 103^. The relevance of topological approaches to this problem lies precisely in their capacity to provide quantitative measure of such observables^57, 104^.

Our findings are relevant in the field of hypnotizability, as they indicate a different embodiment of mental images supporting the socio-cognitive theories of the response to suggestions, and in cognitive science, as the hypnotizability-related information processing can be a model to describe how different selves may emerge from neurophysiological assets. In this respect, the question “whether highs and lows live in the same world”^71^, that is how they represent and reconstruct sensori-motor information, seems to be efficaciously addressed by TDA.

## Methods

### Subjects

Hypnotizability was measured in a sample of 150 healthy students of the University of Pisa through the Italian version of the Stanford Scale of Hypnotic Susceptibility Scale (SSHS), form A^105^ and classified as highly (highs, SHSS score ≥ 8/12), medium (mediums, SHSS score: 5 - 7/12) and low hypnotizable (lows, SHSS score ≤ 4/12). Among them, 20 consecutive Highs (SHSS score (mean+SD): 9.47 ± 1.65) and 20 consecutive lows (SHSS, mean+ SD: 2.01 ± 1.89) with negative anamnesis of neurological and psychiatric disease, drug free for at least the latest 2 weeks were invited to complete the Edinburgh Handedness Inventory (EHI) aimed at characterizing their handedness^106^. Two Highs and 1 Low subject were excluded because not strictly right-handed (EHI score < 16/18); the remaining 18 Highs ( age, 23 ± 1.9 yrs 11 females) and 19 Lows ( age, 22.7 ± 1.5 yrs; 10 females) were invited to participate in the second part of the study consisting of EEG recording during various experimental conditions.

### Experimental Procedure

During the experimental sessions, which were conducted between 11 a.m and 2 p.m., at least 2 hours after the last meal and caffeine containing beverage, participants were comfortably seated in a semi-reclined arm-chair in a sound-and light-attenuated, temperature controlled (21 degree Celsius) room. The experimental procedure was entirely conducted with eyes closed and consisted of 5 trials. Each trial included a basal and a task condition lasting 1 minute each. In basal conditions participants were invited to relax at their best. The tasks consisted of the visual or kinesthetic imagery of a rotated head position performed before (Time 1: v1,k1) or after (Time 2: v2,k2) maintaining an actually rotated head posture (rr). The order of the visual and kinesthetic imagery-which was the same for the trials performed before and after rr - was randomized among subjects. The scripts for the visual and kinesthetic imagery were prepared ad hoc and aimed at presenting the same mental images studied in earlier behavioral experiments^48^. They were read to each participant immediately before the specific condition (v1,v2,k1,k2) and sounded as follows: for visual imagery: *“Please imagine that your head is rotated towards the right side; try to see your chin aligned with your shoulder and maintain this mental imagine until I will tell you to stop”*; for kinesthetic imagery: “ *Please imagine that you head is rotated towards the right side; try to feel the tension of your neck muscles and maintain this mental imagine until I will tell you to stop”*. After each imagery task, subjects were invited to score on scales the vividness of their mental images and the experienced cognitive effort a scale from 0 (minimum) to 10 (maximum). At the end of each imagery condition they were also asked to indicate whether they had obtained better mental image of rotated head at the beginning, at the end or in the middle of the task, in order to allow to select for analysis the 20 EEG seconds in which they reported to have performed the requested task at their best. During the task of actually rotated head position, the subjects were instructed to maintain their head rotated toward the right side in order to align their chin with the right shoulder until the experimenter will allow them to change their head posture. The head position was visually controlled by one of the experimenters sitting behind the subject throughout the session.

### EEG acquisition and processing

EEG was acquired (sample rate:1000 Hz) through a Quick-CapEEG and QuickCell^®^ (Compumedics NeuroMedical Supplies) standard system. Thirty-two EEG electrodes were placed on the scalp according to the 10-20 International System. In addition, 2 auricular (A1, A2), 4 eye (right and left medial/lateral) and 2 EKG electrodes (standard DI lead) were placed. The electrode used as reference during acquisition was FCz; off-line the signal was referred to A1/A2 and FCz was restored. Electrodes impedance was kept under 10 *k*Ω. Filters were applied a posteriori (notch at 50 Hz, bandpass 0.5-45 Hz) for the signal pre-preprocessing. The bad channels interpolated using the spherical interpolation method (EEGLAB `pop_interp` function) were maximum 1 in each subject. Then, data were decomposed into source component using Indipendent Component Analysis (infomax ICA algorithm, EEGLAB function `runica`). Components were visually inspected in order to remove motion, muscular and ocular artefacts. The signal (total duration = 10 min) was divided into 20 seconds epochs (20.000 samples). The automatic selection process (deletion criteria: amplitudes > 100*μ*V, median amplitude > 6SD of the remaining channels) removed not more than 1 epoch in each subject. For basal conditions and for the condition of really rotated head, the earliest less noisy 20 seconds interval (mostly between sec 0 and 30) was chosen for analysis. For the imagery tasks the selected epoch was chosen in each subject according to the interview (between sec 10 and 30 in 69% of Highs, and 64% of Lows).

### Variables and statistical analysis

#### Subjective reports

The scores of the vividness of the mental images (range: 0-10) and of cognitive effort (range: 0-10) associated with the imagery tasks were analyzed through repeated measures ANOVA with Hypnotizability as between subjects factor. Imagery modality (visual, kinesthetic) and Time (T1,before rr - real rotation (v1,k1); T2, after rr (v2, k2)) were within subjects factors.

#### EEG homological signal analysis

Persistent homology on a transformation on the correlation matrix obtained from EEG temporal series by subject, group and condition. Persistence diagrams are the main feature obtained from persistent homology analysis. We have used them joined to a similarity measure adapted to persistent diagrams to differentiate groups and conditions. The EEG topological parameters (persistence diagrams) were analysed through a measure of similarity adapted to this feature which depends on a free parameter understood as the desired scale of comparison. We define a normalized measure of similarity between two persistent diagrams based on Reininghaus kernel^68^. We first computed similarity between condition and basal mode for each subject inside the same group (e.g. highs v1 condition vs basal condition) and we showed the mean similarity inside the group and compared it between groups (Highs and Lows) for different conditions. This analysis was followed by an uniformity analysis intra group. Secondly, we computed similarity between real rotation and conditions (e.g. condition v1 and rr) on the same subject and compare results between groups (Highs and Lows). All these results are showed with their corresponding confidence interval at 95% (assuming a Gaussian error distribution).

### Topological markers: persistent homology and persistence diagrams

One of the main tools in TDA is persistent homology (PH)^107, 108^. Persistent homology captures topological data information, where topological in this context refers to the shape of the data. In other words, persistent homology shows the evolution of “holes” that define the shape of the data at different dimensions (0-dimensional holes correspond to the set of connected components (points), 1-dimensional holes correspond to cycles, 2-dimensional holes correspond to cavities, and so on). PH provides a robust multiscale summary of a whole dataset thanks to its capacity to capture the global topological structure of point cloud data or a weighted network rather than their local geometric behavior, thereby making the summaries robust to missing data and small errors. One of its big potential is its threshold freeness. While homology provides information about the existence of n-dimensional holes at a given threshold, PH use all possible thresholds to follow the evolution of each n-dimensional hole. The picture of possible subdata sets for each threshold is called filtration. More details on persistent homology and it mathematical definition are provided in the SI.

PH accepts as input point cloud data and weighted networks. In our study we work with weighted graphs obtained from correlation matrices computed on EEG time series signals. A weighted network is a way to represent pairwise relationships and encode their strength. In our case, the nodes “EEG points: electrodes” and links between nodes are weighted by the Pearson correlation between the nodes’ signals. In order to consider a filtration from stronger to weaker correlations, we transform the correlation values by using one minus the absolute Pearson’s correlation coefficient. We then convert a weighted network to a series of simplicial complexes by taking the clique complex of progressively increasing thresholded networks.

A clique is a subset of vertices such that they induce a complete subgraph (K_i_ is the full connected graph of i nodes). That is, every two distinct nodes in the clique are adjacent. A simplicial complex is a topological space constructed by simplices where simplices are points, lines, triangles, and their n-dimensional counterparts (see Figure S.2 in SI). We obtain a simplicial complex from a network through cliques by mapping k-cliques to (k-1)-simplices (see Figure S.2 in SI). Interestingly, although all the information is encoded in the pairwise interactions, converting a graph to a simplicial complex reveals mesoscopic organizational structure that was not appreciable at the network level, thanks to the non-locality of the topological invariants of the simplicial complex. In the presented work, the filtration used for the computation of persistent homology consists in a family of clique complexes, *X*_*i*_ one inside the other ( ⋯ ⊆ *X*_*i*−1_ ⊆ *X*_*i*_ ⊆ *X*_*i*+1_ ⊆ …) obtained from the progressive thresholding of a weighted network as in Figure S.3 in the SI shows.

Persistent homology can be computed to find n-dimensional holes. Here we focus on 1-dimensional holes that correspond to cycles which correspond to the generators of the homology group *H*_1_. Roughly speaking a hole is defined as a closed object which is not a boundary of any other object. That is, a n-dimensional hole encloses a subspace, a cavity. Thereby, PH consist in computing *H*_1_(*X*_*i*_) for each *X*_*i*_ along filtration.

The output of PH is summarized in a topological invariant called persistence diagram. A persistence diagram consist of a set of tuples of numbers, corresponding to the starting point (birth step) and ending point (death step) of each hole along the filtration. The importance of a hole is proportional to the length of its interval of existence, called its persistence. In other words, cycles that live for large intervals in the filtration capture the main features of the shape of the dataset, while shorter ones are usually considered as noise. In terms of persistence diagrams, this interpretation gives more importance to points further from the diagonal.

In order to compare persistent diagrams we use a similarity measure defined by a kernel function (see example in Figure S.4 in SI) based on^68^. Given two persistence diagrams *F* and *G* (sets of birth and death cycle points), we define: 
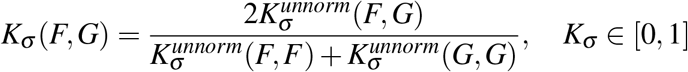
where, 
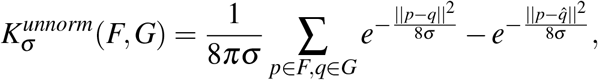

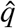 is the mirror point of *q* (if *q* = (*q*_1_, *q*_2_), 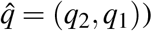) and *σ* is a parameter related to the filtration step size. Note that *K*_*σ*_ (*F*, *F*) = 1 for any *F* persistence diagram. similar the closer their *K*_σ_ is to 1.

## Acknowledgements (not compulsory)

EIM and GP are supported by the ADnD project of Compagnia San Paolo.

## Author contributions statement

ELS designed the experiment, EIM and GP designed the analytical framework, LC and ELS conducted the experiment and preprocessed the data, EIM,GP and AP analysed the results. All authors wrote and reviewed the manuscript.

## Mathematical formulation of homology and persistent homology

We introduce some technical notions in the context of persistent homology: chain complex, boundary map and the formal definition of homology^1, 2^.

The set of *n*-dimensional chains *C*_*n*_(*X*) is the formal sums of *n*-simplices. of a simplicial complex *X* Formally written as, 
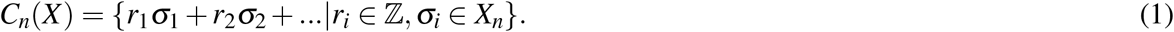

Then we can define a *boundary map* ∂_*n*_ between *n*-dimensional chains *C*_*n*_(*X*) to (*n* − 1)-dimensional chains *C*_*n* − 1_(*X*) that corresponds to our intuitive notion of boundary.

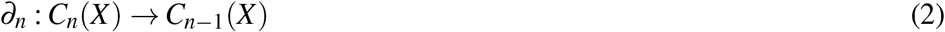

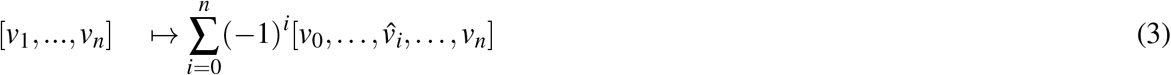
where the hat denotes the omission of the vertex and this map satisfies ∂_*n*_∂_*n*+1_ = 0 ∀*n*. For example, if we have a simplex of dimension 2 (a full triangle), it will be converted to its boundary, that is, three concatenated edges (simplices of dimension 1). The boundary of a simplex is the alternating sum of restrictions to its faces.

A simplicial complex *X* induces the *chain complex*, ⋯ → *C*_*n*+1_ → *C*_*n*_ → *C*_*n*−1_ → … through boundary maps …∂_*n*+2_, ∂_*n*+1_, ∂_*n*_, ∂_*n*−1_,…

The *n-homology* of this complex is defined by the quotient of two vector spaces, the kernel of the map ∂_*n*_ quotiented by the image of the boundary map one upper dimension, ∂_*n*+1_, 
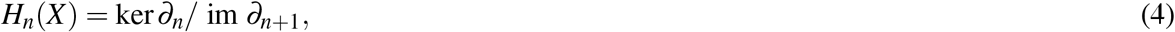
where *n* indicates the dimension of the generators in the homology group. We call the kernel ker ∂_n_ the *n*th cycle module, denoted *Z*_*n*_, and the image im ∂_*n*_ the *n*th boundary module, denoted *B*_*n*_.

Formally, given a simplicial complex *X*, a filtration is a totally ordered set of subcomplexes *X*_*i*_ ⊂ *X*, that starts with the empty complex and ends with the complete complex, indexed by the nonnegative integers, 
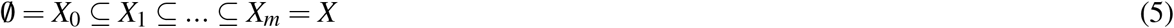
such that if *i* ≤ *j* then *X*_*i*_ ⊆ *X*_*j*_. The total ordering itself is called a filtration. The subcomplexes *X*_*i*_ are the analog of the sublevel sets in the Morse function setting^3^.

In order to define *persistent homology*^4, 5^, we use superscripts to denote the index in a filtration. The *i*th simplicial complex *X*_*i*_ in a filtration gives rise to its own chain complex 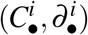 and the *k*th chain, cycle, boundary and homology modules are denoted by 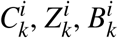 and 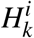, respectively.

For a positive integer *p*, the *p-persistent kth homology* module of *X*_*i*_ is 
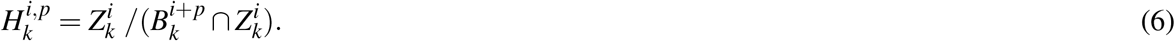

The form of 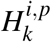 should seem similar to the formula for 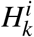, except that instead of characterizing the *k*-cycles in *X*_*i*_ that do not come from a (*k* + 1)-chain in *X*_*i*_, it characterizes the *k*-cycles in the *X*_*i*_ subcomplex that are not the boundary of any (*k* + 1)-chain from the larger complex *X*_*i*+*p*_. Put another way, 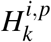 characterizes the *k*-dimensional holes in *X*_*i*+*p*_ created by the subcomplex *X*_*i*_. These holes exist for all complexes *X*_*j*_ in the filtration with index *i* ≤ *j* ≤ *i* + *p*.

## TABLES

**Table S.1.** t-test results on the mean value for different conditions in Figure 2.

| Condition | t-test | p-value |
| --- | --- | --- |
| k1 | 22.75 | 4.83e-20 |
| k2 | 27.00 | 4.19e-22 |
| v1 | 24.86 | 4.22e-21 |
| v2 | 19.60 | 2.85e-18 |
| $\Delta k$ | 65.76 | 4.07e-33 |
| $\Delta v$ | 19.42 | 3.64e-18 |

## Figures

**Figure S.1.**
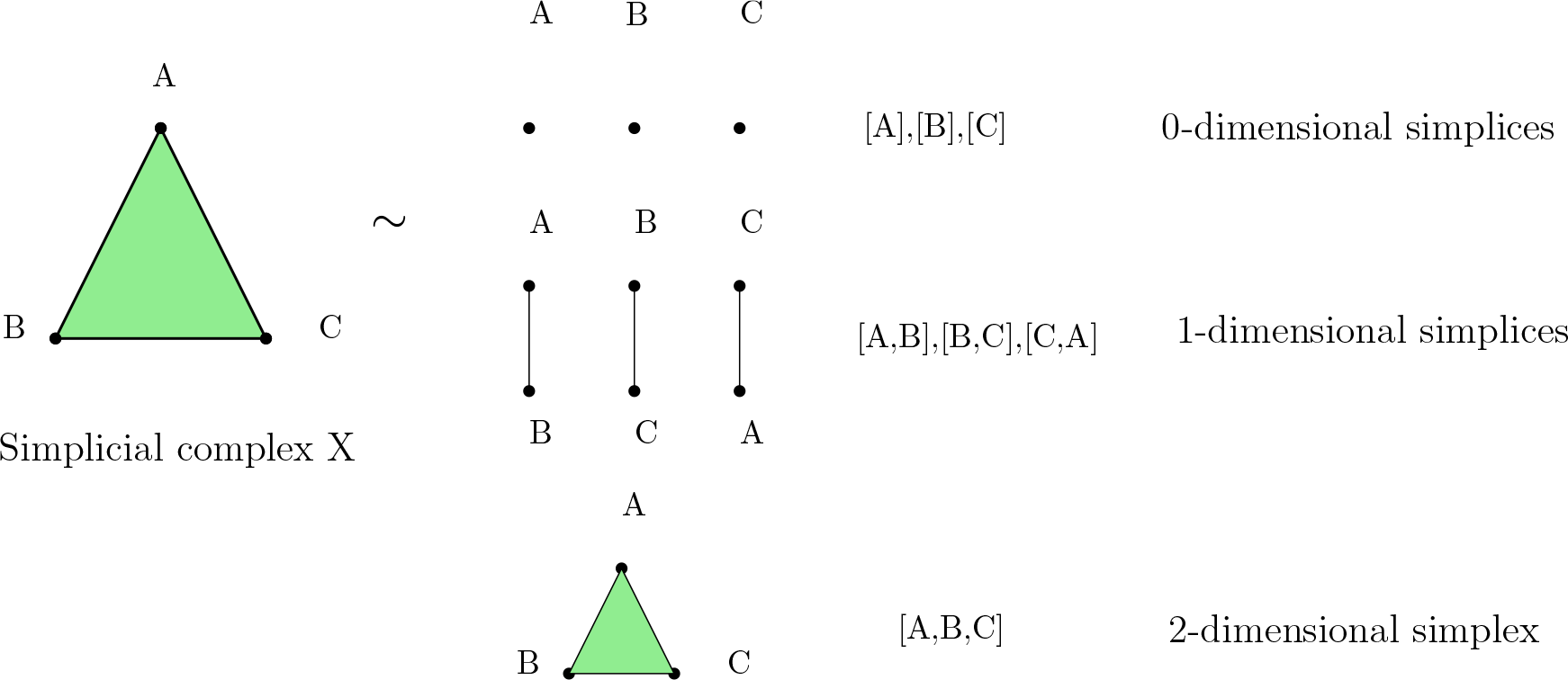
An example of simplicial complex, *X* and its pieces: *X* = {[*A*], [*B*], [*C*], [*A*, *B*], [*B*,*C*], [*C*, *A*], [*A*, *B*,*C*]}. In other words, a full triangle is composed by three 0-simplices (points), three 1-simplices (segments) and a 2-simplex (a full triangle).

**Figure S.2.**
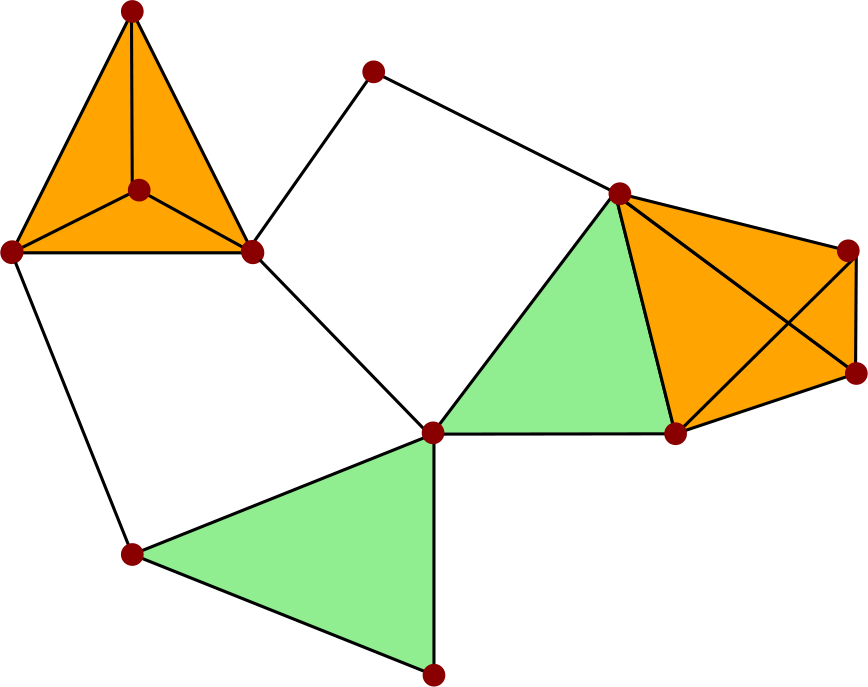
Clique complex of a network. Cliques in the network are mapped (*filled*) to simplices of the corresponding dimension in order to obtain a simplicial complex.

**Figure S.3.**
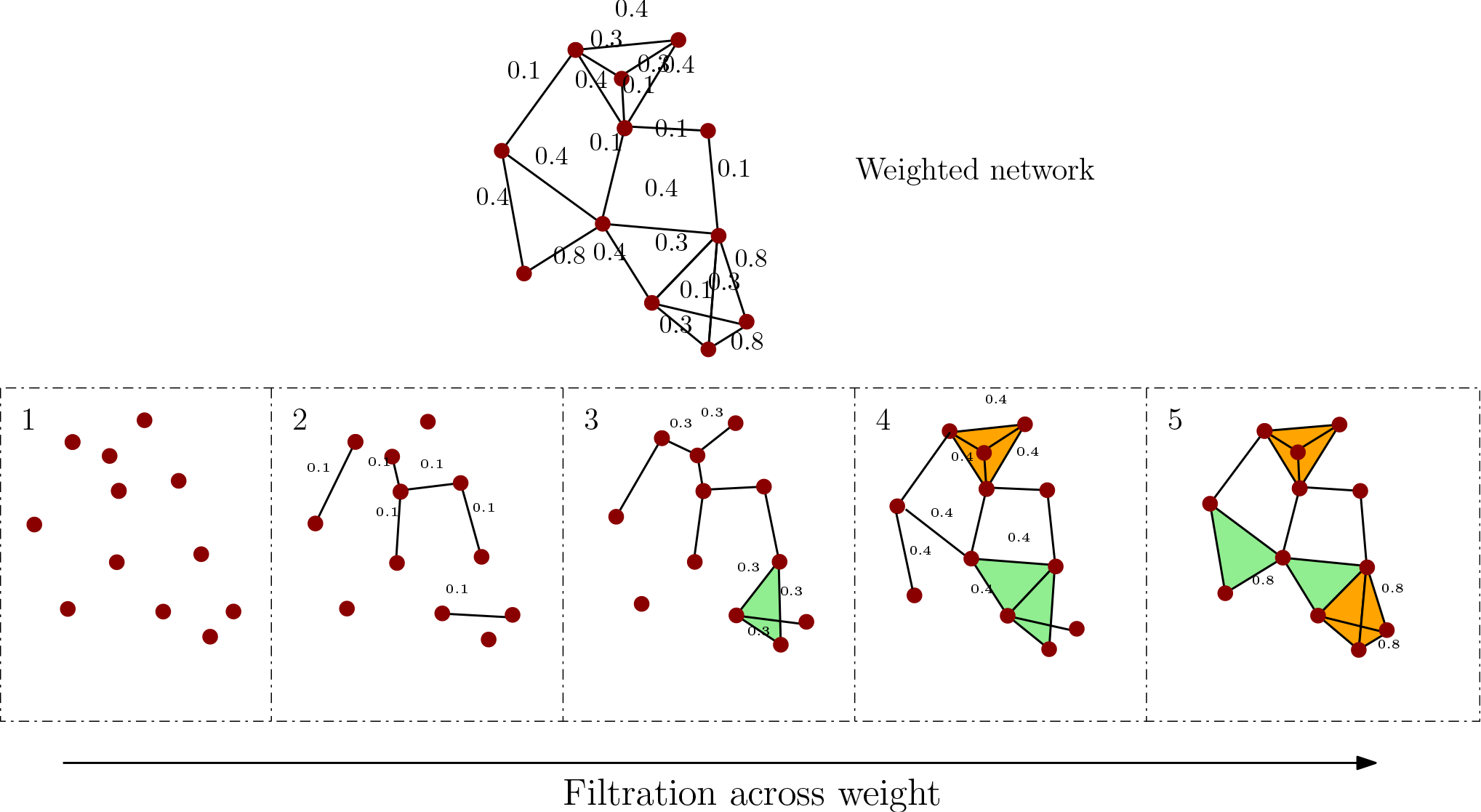
Clique complex filtration over weights of a weighted network.

**Figure S.4.**
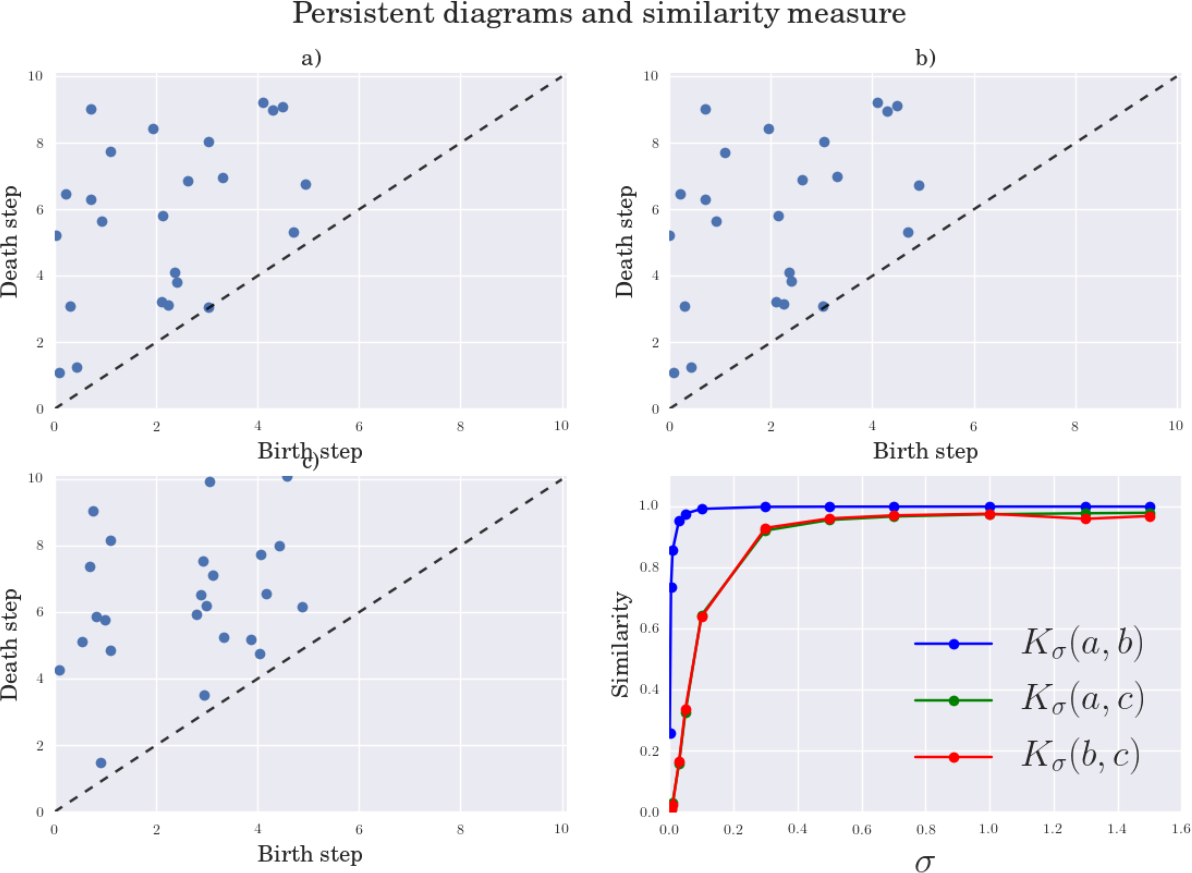
Examples of persistent diagrams in a),b) c) and similarity measure between them. Points further from the diagonal (dotted line) in persistent diagrams represent cycles with a larger persistence along filtration. Similarity between a) and b) is higher than similarity between a) and c) and b) and c), *K*_σ_ (*a*, *c*) ⪡ *K*_σ_ (*a*, *b*) and *K*_σ_ (*b*, *c*) ⪡ *K*_σ_ (*a*, *b*).

**Figure S.5.**
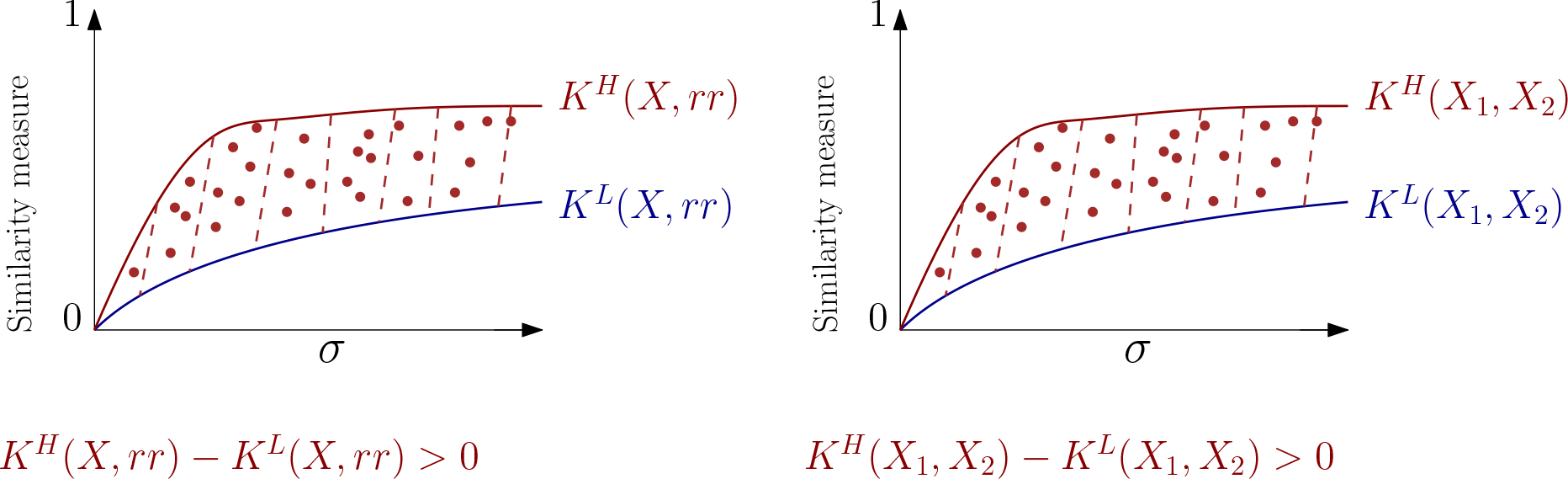
Difference between Highs and Lows of imagery tasks and real rotation across similarity curves obtained from persistence diagrams. Left: Difference between Highs and Lows of imagery tasks alteration of topology relate the actual rotations (*rr*). Positive results shows how Highs are always topological nearer to the real rotation than Lows for all imagery conditions. Right: Difference between Highs and Lows of imagery tasks alteration of topology before (k1,v1) and after (k2,v2) the actual rotation. Positive results show how Highs both in kinesthetic and visual imaginations are always topological nearer to the real rotation or that real rotation does not inference in their imagination than Lows for all imagery conditions.

**Figure S.6.**
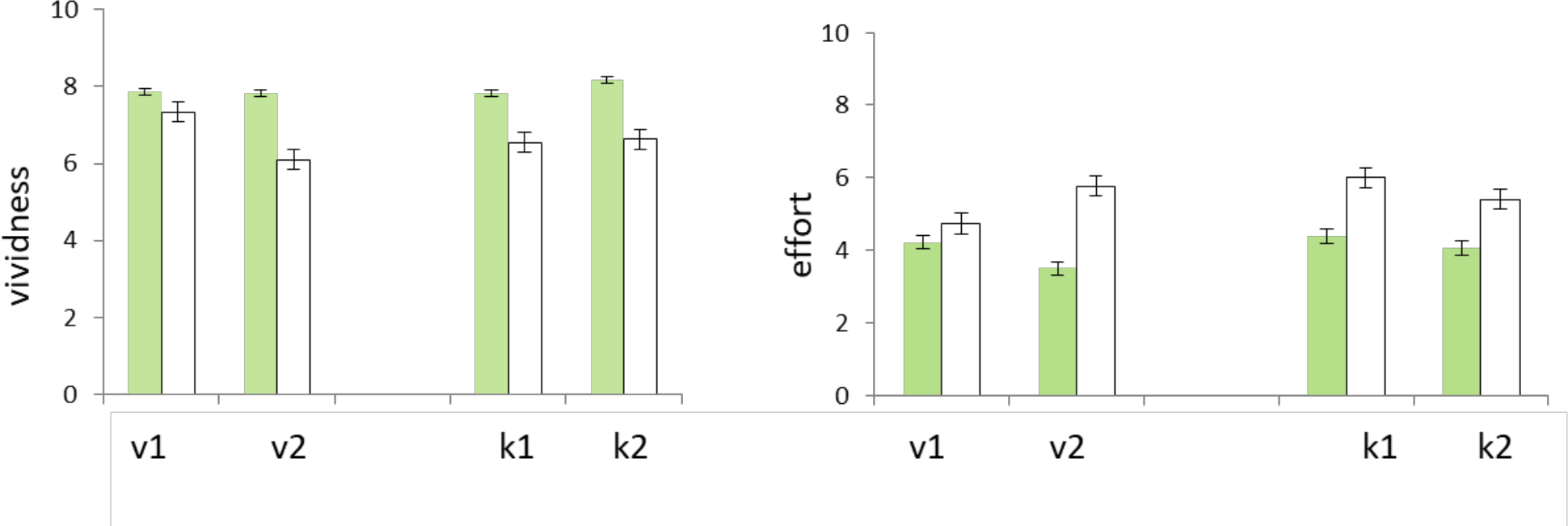
Subjective reports (Mean, SEM) of vividness of imagery (left panel) and cognitive effort (right panel). Highs (green) reported significantly higher vividness and lower effort than Lows. Lows reported lower vividness for later tasks. The Highs’ effort was greater for earlier than for later tasks. v1/k1 and v2/k2: visual and kinesthetic imagery before/after the actually rotated position of the head, respectively. For statistics see text.

**Figure S.7.**
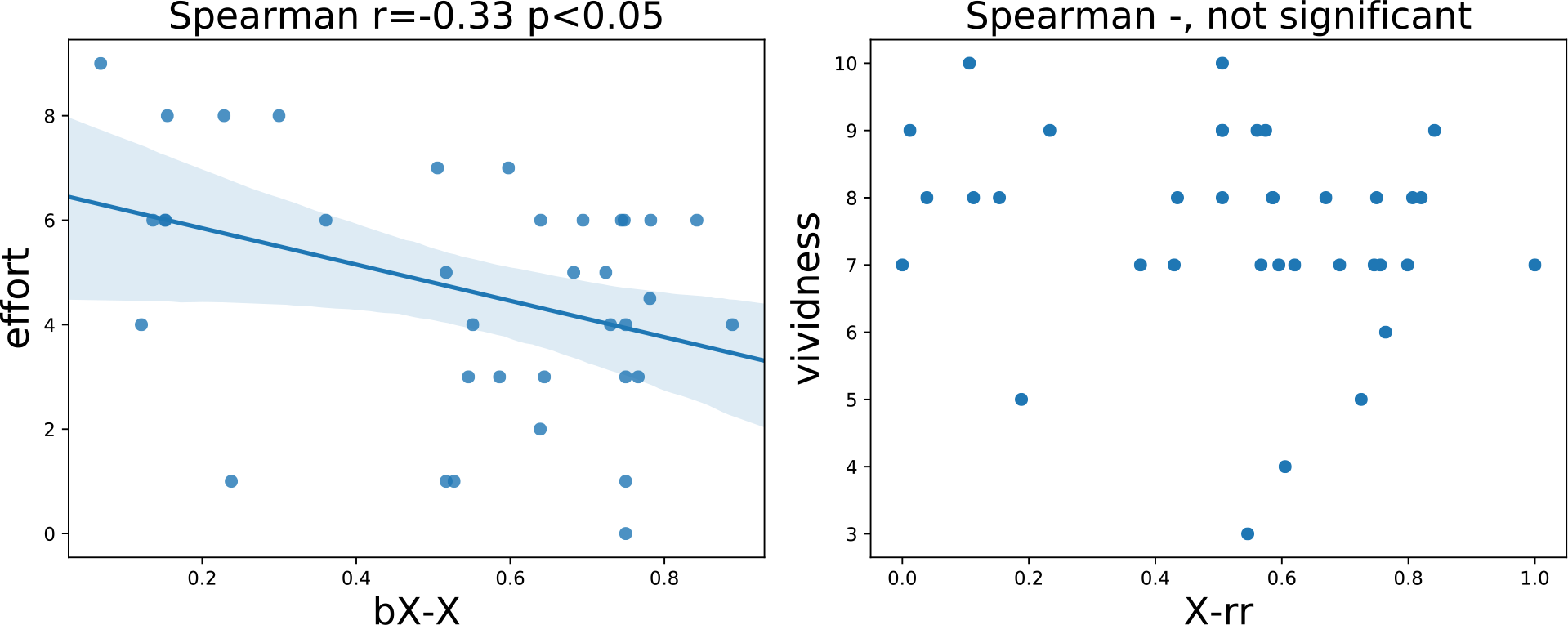
Individual correlations. Aggregating all imagery tasks, we show the relation between the subjective values between effort and Ω(*bX*, *X*) (left), and between vividness and Ω(*rr*, *X*) (right). In the first case, as expected we recover a (significant, *r* = −.33, *p* < .05) negative correlation between basal deviation and reported effort. For vividness at the subjective level our results are inconclusive as we do not find significant correlations.

